# Systemic infection with highly pathogenic H5N8 of avian origin produces encephalitis and mortality in wild mammals at a UK rehabilitation centre

**DOI:** 10.1101/2021.05.26.445666

**Authors:** Tobias Floyd, Ashley C. Banyard, Fabian Z. X. Lean, Alexander M. P. Byrne, Edward Fullick, Elliot Whittard, Benjamin C. Mollett, Steve Bexton, Vanessa Swinson, Michele Macrelli, Nicola S. Lewis, Scott M. Reid, Alejandro Núñez, J. Paul Duff, Rowena Hansen, Ian H. Brown

**Affiliations:** Animal and Plant Health Agency (APHA), Weybridge, Surrey, United Kingdom; APHA Thirsk, North Yorkshire, United Kingdom; Royal Society for Prevention of Cruelty to Animals (RSPCA), East Winch, Norfolk, United Kingdom; APHA Bury St Edmunds, Suffolk, United Kingdom; Department of Pathobiology and Population Sciences, Royal Veterinary College, North Mymms, Hertfordshire, United Kingdom; APHA Diseases of Wildlife Scheme (DoWS), Penrith, Cumbria, United Kingdom

## Abstract

Europe has experienced extensive outbreaks of highly pathogenic avian influenza (HPAI) during the autumn/winter 2020/21 season. These avian influenza A viruses are highly transmissible and have infected over 1000 commercial and backyard poultry premises in Europe in this period causing high mortality. The impact on wild bird populations has also been significant, with over 400 detections in at least 47 different species reported across Europe as being positive with the H5N8 virus. Although different H5Nx combinations within the H5 clade 2.3.4.4b have been detected, the H5N8 subtype has predominated both in wild birds and domestic poultry outbreaks. In the UK there have been 22 outbreaks of H5N8 in domestic poultry and captive birds and more than 300 wild bird detections involving H5N8 over the autumn/winter 2020/21 period to April 2021. Here we detail the series of events surrounding the detection of an H5N8 influenza A virus of avian origin in five swans, a fox and three seals in a wildlife rehabilitation centre.

## Outbreak detection

An episode of unusual disease and mortality at a wildlife rehabilitation centre led to the detection of H5N8 influenza A virus of avian origin in five swans, a fox and three seals (Figure 1). Four juvenile common seals (*Phoca vitulina*), one juvenile grey seal (*Halichoerus grypus*) and one juvenile red fox (*Vulpes vulpes*) died or were euthanased over a two-day period. The fox had died suddenly after a short period of non-specific malaise and inappetence. The seals had exhibited sudden-onset neurological signs, including seizures prior to death or euthanasia.

**Figure 1:**
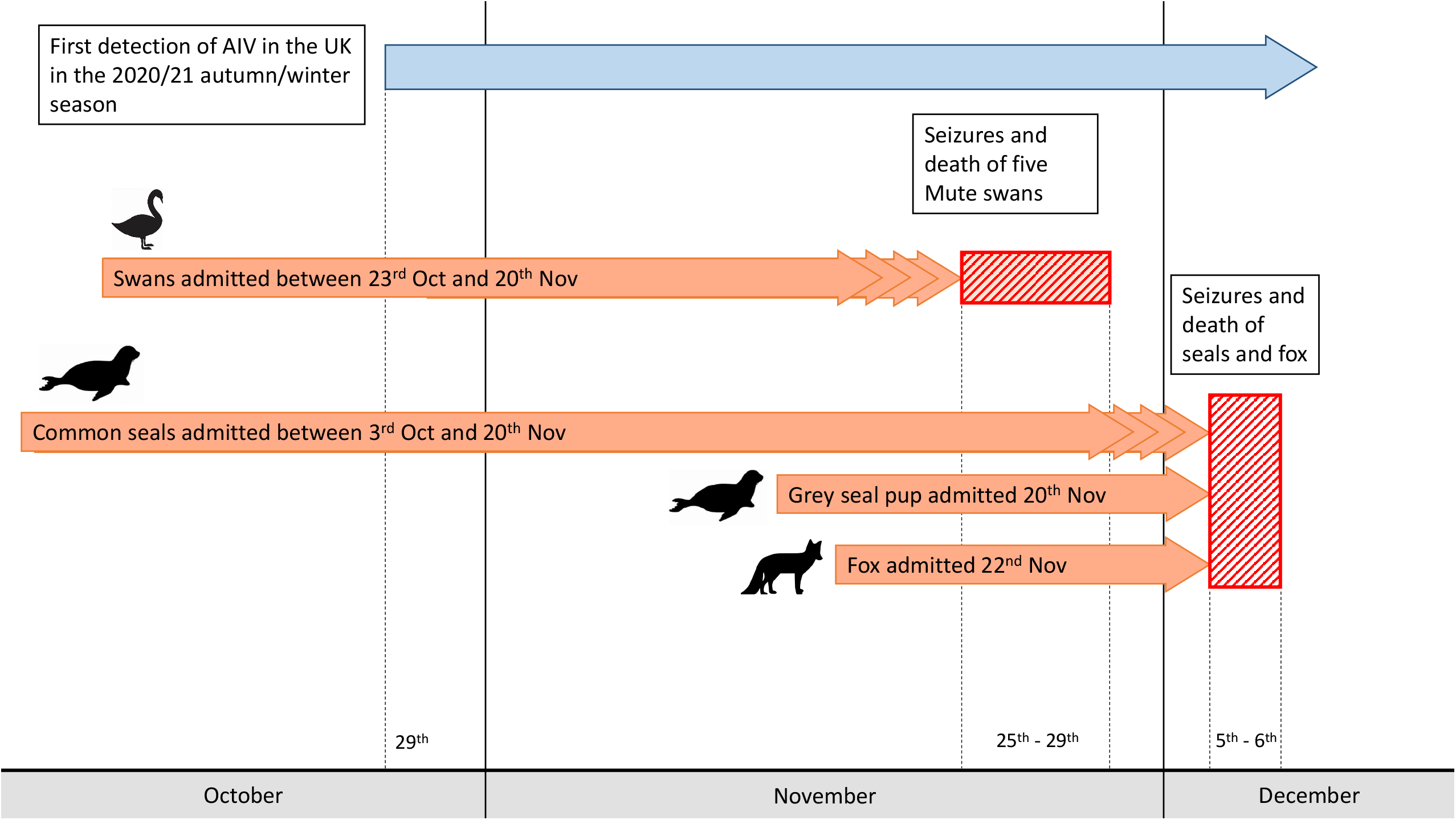
Timeline of the disease event.

This episode occurred approximately one week after the deaths or euthanasia of five mute swans (*Cygnus olor*) held in isolation at the centre, following acute onset malaise and terminal seizures. The five swans were submitted for examination and testing under the Avian Influenza Wild Bird Surveillance Scheme [1] and tested positive for highly pathogenic avian influenza virus (HPAIV) of the H5N8 subtype.

The unusual spatio-temporal cluster of unexplained mortality and neurological disease in multiple avian and non-avian species warranted further investigation. The fox and five seal carcases were submitted to an Animal and Plant Health Agency (APHA) regional Veterinary Investigation Centre for routine examination under the APHA Diseases of Wildlife Scheme with a view to achieving an aetiological diagnosis. Influenza of avian origin was not suspected and none of the captive birds at the centre showed any clinical signs of disease.

## Methods

### Post-mortem examination, tissue sampling and histopathological investigation

A thorough clinical history was collected from the rehabilitation centre. The carcases of three of the five swans, the fox and three of the five seals (one grey seal and two common seals) were subject to full post-mortem examination (PME). Two swans were submitted frozen and did not undergo PME – but oral and cloacal swabs were taken from these carcases for virological testing. Two common seal carcasses were deemed too autolysed for examination and not sampled.

Samples of heart, lung, liver, kidney, spleen and brain tissues were taken from the swan, fox and seals, fixed in 10% neutral buffered formalin and processed for Haematoxylin and Eosin staining and immunohistochemistry (IHC). IHC was undertaken using anti-influenza A nucleoprotein primary antibody (Statens Serum Institute, Denmark), as previously described [2], and anti-canine distemper virus nucleoprotein mouse monoclonal primary antibody (Biorad) in selected fox tissues.

Plain swabs were taken from the oropharynx and cloaca of each of the swans and similarly from the nasal cavity and rectum of each of the three seals examined for preliminary influenza virological testing. Tissue samples of brain, liver, kidney, spleen and lung were taken from both the seals and the fox and were stored at −80°C until required.

### Virological investigation

Swabs taken from the swans and seals and tissue samples taken from the seals and fox were assessed for influenza A nucleic acid using a screening real time reverse-transcriptase polymerase chain reaction (rtRT-PCR) assay followed by subtype specific rtRT-PCRs. Where H5 subtype specific PCR detected positive samples, the haemagglutinin cleavage-site sequence (CS) was determined (all protocols available on https://science.vla.gov.uk/fluglobalnet/).

Virus isolation was undertaken on PCR positive samples from the swans, seals and fox using 9 to 11 day-old specified pathogen free embryonated fowls’ eggs [3, 4]. Viral isolates obtained were then used to generate whole-genome sequence (WGS) data using an Illumina MiSeq [5]. For comparative genetic analysis, recent H5 2.3.4.4b virus HA sequences were downloaded from the GISAID EpiFlu database (https://platform.gisaid.org/ accessed on 17th March 2021). WGS were deposited on the GISAID database under accession numbers: EPI_ISL_1123360, EPI_ISL_2081527 and EPI_ISL_2081528.

Additionally, samples obtained from the fox were subjected to RNA extraction and rtRT-PCR to evaluate the presence or absence of rabies virus and canine distemper virus nucleic acid [6, 7] and rtRT-PCR assays for detection of *Leptospira* [8] were carried out on kidney tissue from each of the three examined seals. Additionally, RNA obtained from the fox samples were subjected to sequence-independent single-primer amplification (SISPA) [9] to generate double-stranded cDNA that was then sequenced using an Illumina MiSeq. The WGS data obtained from these samples was then used to exclude other viral agents. Firstly, reads that aligned to the *Vulpes vulpes* genome were removed using Burrows-Wheeler Aligner [10] and Samtools [11]. The non-host reads were then *de novo* assembled to produce contiguous sequences using SPAdes [12] which were then used to screen custom viral databases obtained from ViPR [13] using BLAST+. For this, databases of the *Bornaviridae, Circoviridae, Flaviviridae, Herpesviridae, Paramyxoviridae, Parvoviridae, Pheuiviridae* and *Rhabdoviridae* families were used.

## Results

### Clinical Setting

The swans, grey seal and fox were housed within the isolation unit of the centre. The isolation unit comprised 17 individual cubicles accessed by a central corridor. Animals in separate cubicles did not have direct contact. The unit had been purposefully designed to reduce the potential for transmission of infection between cubicles and biosecurity measures were in place, such as protective clothing and decontamination steps between cubicles.

The five juvenile swans were rescued from different locations and brought to the centre for treatment between the 23^rd^ of October and the 20^th^ of November. These birds were admitted for various reasons, including trauma and being underweight and weak. The swans were housed indoors within the centre’s isolation unit, mixed in groups of up to four with other rescued adult swans. Each of the swans had been recovering uneventfully until the sudden onset of lethargy and death or euthanasia between the 25^th^ and 29^th^ of November (Figure 1). Clinical signs were not observed in the remaining adult swans within the isolation unit. The isolation unit also contact several juvenile mallard ducks (*Anas platyrhynchos*) that had been brought to the centre as ducklings two-to-three months prior. There was no clinical disease or mortality recorded in these birds.

The four common seals were estimated to be aged between five and six months-old and had, individually, arrived at the facility between one and two months prior to the disease episode (Figure 1). They had been admitted for various reasons, including poor body condition, superficial bite wounds and lungworm. In each case the animals had been responding well to treatment and supportive care up until the sudden onset of seizures and subsequent death or euthanasia. All four common seals that died had been present in the isolation unit at the time the swans were affected – occupying cubicles directly opposite those of the swans – but had subsequently been moved into another area of the facility. these seals had had close contact with other common seals in other areas of the facility, none of which became ill.

The grey seal was a two-week-old pup admitted for care following maternal abandonment two weeks prior (Figure 1). It was housed in the isolation unit in a cubicle opposite the swans. The seal was in good body condition and deemed to be progressing well until the sudden onset of clinical signs – including pyrexia, facial twitching and stupor – whereupon it was euthanased on welfare grounds.

The fox had been in the isolation unit of the centre for two weeks (Figure 1). It had presented to the facility with large areas of alopecia and skin crusts over the body and limbs, consistent with mange. It had been receiving treatment and was reported to be progressing well until the sudden onset of malaise and inappetence and was found dead the following morning.

### Pathological Investigation

The body condition of the three swans examined ranged from good to poor. Gross findings among the carcases included petechiae in the liver and epicardium and opacity of the air sacs. Microscopic examination of tissues from three birds revealed multifocal, necrotising, non-suppurative myocarditis, hepatitis, splenitis, nephritis and encephalitis, along with intralesional presence of influenza A virus antigen by IHC.

The body condition of the three examined seals was judged to be fair. Gross examination of two common seals revealed generalised lymphadenomegaly and multiple pale foci in the lungs. One common seal also showed congested meninges. The grey seal pup showed generalised lymphadenomegaly but no other gross changes. Microscopic examination of the seal tissues revealed mild, eosinophilic, interstitial pneumonia in the two common seals – consistent with lungworm infection – and severe, necrotising, non-suppurative polioencephalitis in both common seals and the grey seal. The lymph node sections showed non-specific reactive hyperplasia, accounting for the gross enlargement of these organs. Immunohistochemistry for influenza A virus revealed multifocal immunolabelling in the neurons within the grey matter of the brain in all three seals, in close association with the inflammatory lesions (Figure 2a). Virus antigen was, however, absent in the lung, liver, kidney and lymph node of the seals.

**Figure 2:**
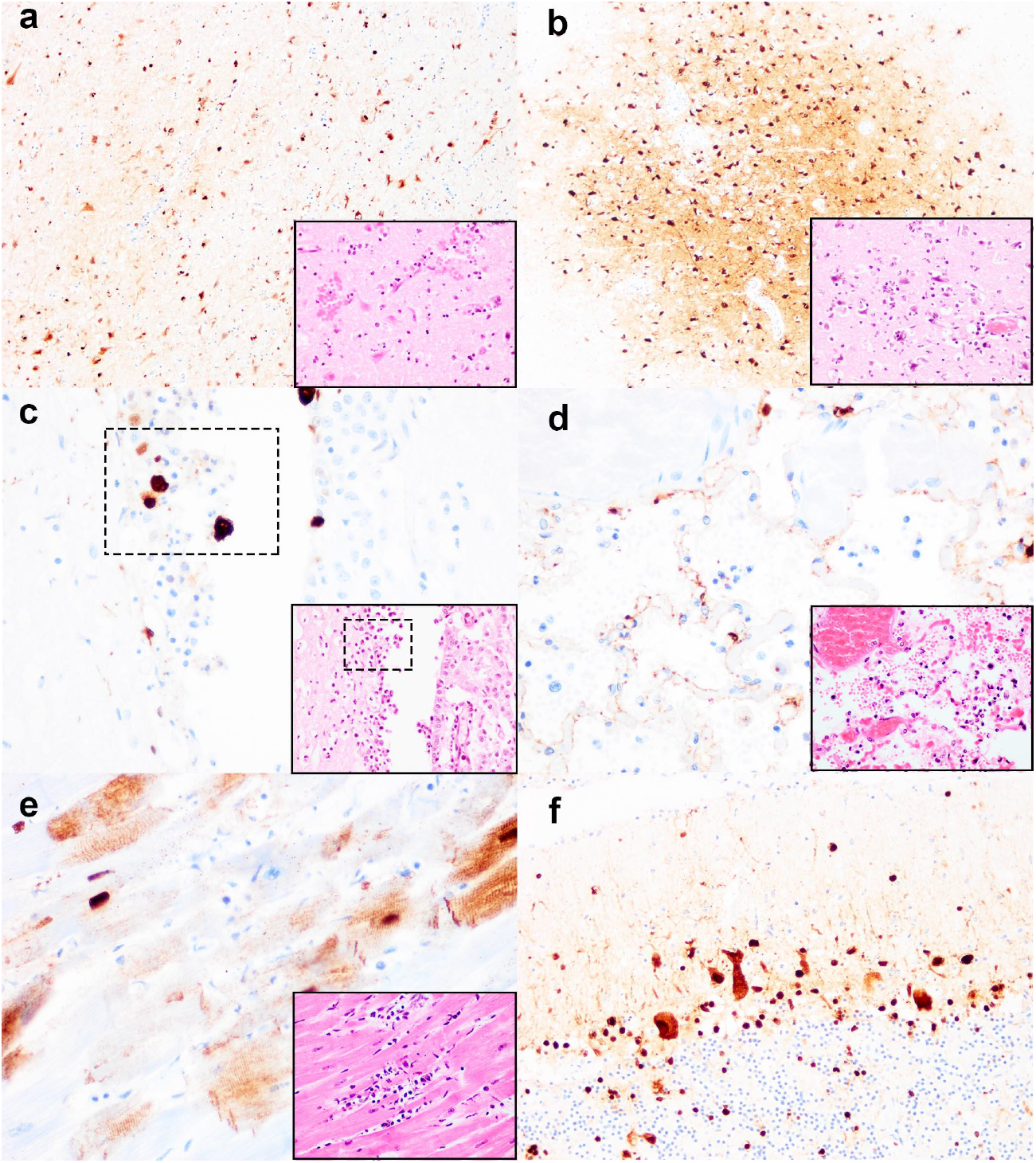
Histopathology and immunohistochemistry of the grey seal (Halichoerus grypus) and red fox (Vulpes vulpes) infected with HPAI H5N8. (a) Non-suppurative polioencephalitis and presence of virus antigens in neurons in the cerebrum, common seal. (b) Non-suppurative polioencephalitis with neuronophagia and association of virus antigens, fox. (c) Ependymal necrosis (inset, box=region of interest) and the association of virus antigens, fox. (d) Diffuse alveolar damage and presence of virus in type I alveolar pneumocytes, fox. (e) Cardiomyonecrosis associated with virus antigens in cardiomyocytes, fox. (f) Virus antigens in granular and molecular layer of the cerebellum, fox. Original magnification 10x (b, d), 20x (f), 40x (c, d, e and insets). Serial tissue sections were stained with haematoxylin and eosin, and for influenza A nucleoprotein using IHC. Insets show histopathology.

Gross examination of the fox revealed large areas of alopecia and crusts affecting the skin of the body and limbs, mild splenomegaly and generalised reddening of the lungs. Microscopic examination of the brain, lung and heart revealed a severe, acute, non-suppurative, polioencephalitis (Figure 2b and ventriculitis (Figure 2c); severe, acute, necrotising, non-suppurative interstitial pneumonia (Figure 2d); and a mild, acute, non-suppurative myocarditis (Figure 2e). Within these lesions, IHC confirmed the presence of influenza virus antigens amongst neurons in the cerebrum, (Figure 2b) and cerebellum (Figure 2f), ependymal cells (Figure 2c), alveolar type I pneumocytes (Figure 2d) and cardiomyocytes (Figure 2e). IHC for canine distemper virus performed on sections of brain and lung was negative.

### Virological assessment

Oropharyngeal and cloacal swabs from all five swans tested positive for H5N8 (Table 1a). Whole genome sequence data was generated for one of the swans submitted (A/mute swan/England/234135/H5N8 2020/12/01)

**Table 1:**
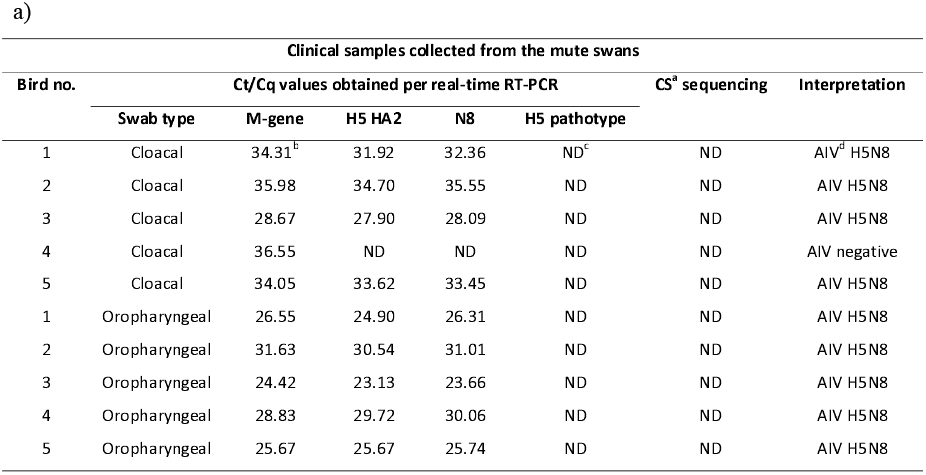
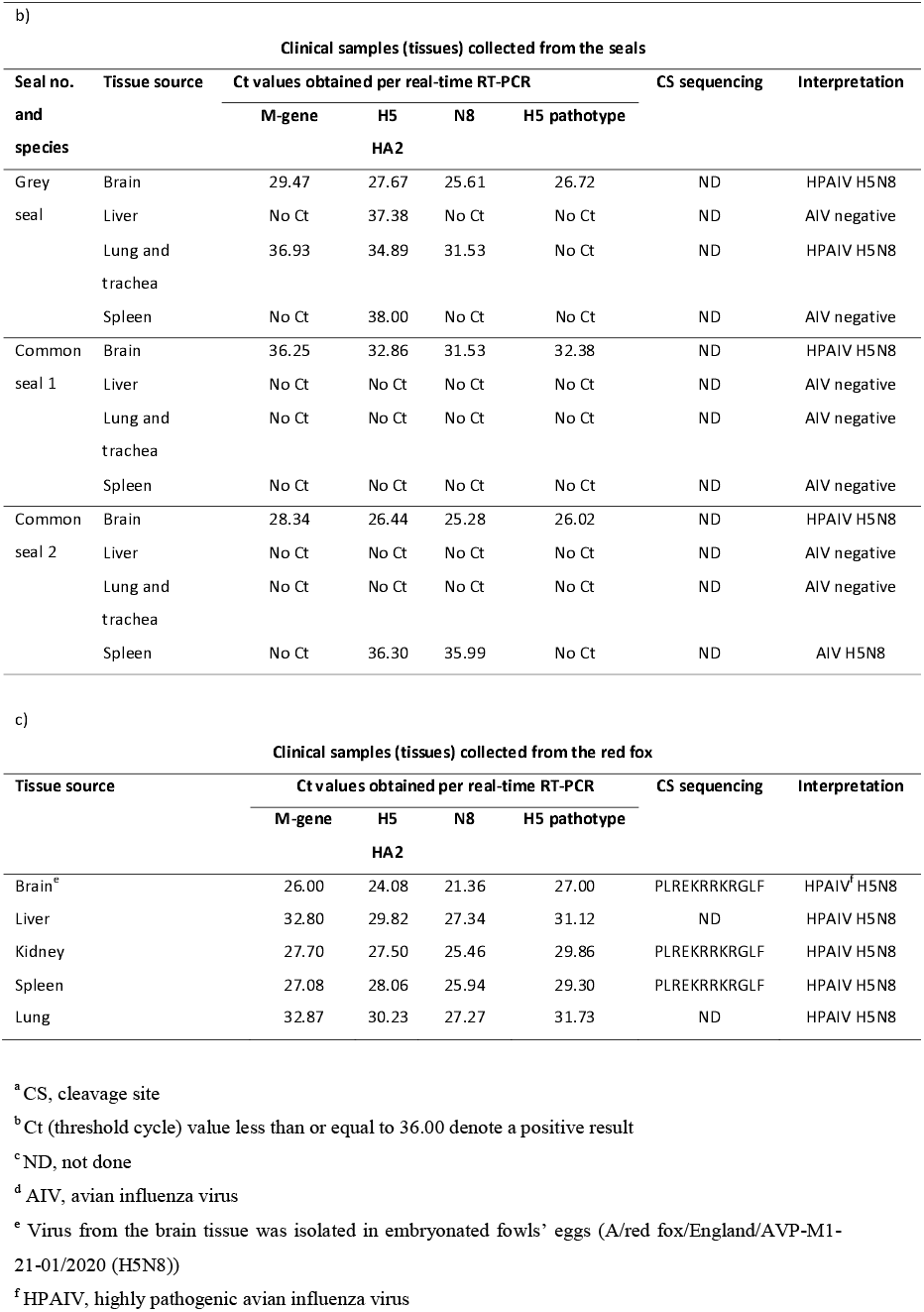
Results from real-time RT-PCR and virus isolation testing of clinical material collected from: a) the Mute swans; b) the fox; and c) and d) the seals

Nasal and rectal swabs taken during PME of the three seals tested negative by PCR for influenza A virus. However, influenza A virus (H5N8) nucleic acid was detected in the brain and lung of the grey seal and the brain of two common seals (Table 1b). Virus isolation was undertaken on pooled samples with successful isolation from the pooled brain tissue of seals (A/seal/England/AVP-031141/2020 (H5N8)).

Influenza A viral RNA was demonstrated in each of the fox samples (brain, liver, kidney, spleen and lung) with PCR subtyping determining the influenza A viral subtype as H5N8 (Table 1c). Virus was isolated from the fox brain tissue (A/red fox/England/AVP-M1-21-01/2020 (H5N8)).

The HA cleavage site (CS) motif from selected fox and seal samples had the amino acid sequence PLREKRRKRGLF, consistent with 99% of CSs characterised across avian H5N8 HPAI viral sequences from the UK during autumn/winter 2020/21.

Analysis of the whole genome sequence (WGS) data generated from the fox, seal and swan samples demonstrated high sequence similarity (>99.9% nucleotide identity across all gene segments). Alignment of the viral haemagglutinin (HA) and neuraminidase (NA) genes demonstrated that virus from both the fox and seals clustered closely with the viruses detected in the swans from the same wildlife centre (Figure 3a (HA) and 3b (NA)). Comparison of WGS data from the fox, seal and swan identified a total of 33 amino acid substitutions associated with altered virulence in the literature that were common amongst the mammalian derived sequences and that have been identified in the majority of avian influenza A virus H5N8 sequences generated during the current epizootic (Figure 3c). The genomes of the fox, seal and swan-derived viruses were homologous at the amino acid level with the exception of a single amino acid substitution at position 701 in the PB2 protein (D701N) that was only present in all of the fox and seal sequences. Analysis of WGS data obtained from the fox samples demonstrated a lack of other viral agents, including canine distemper virus.

**Figure 3.**
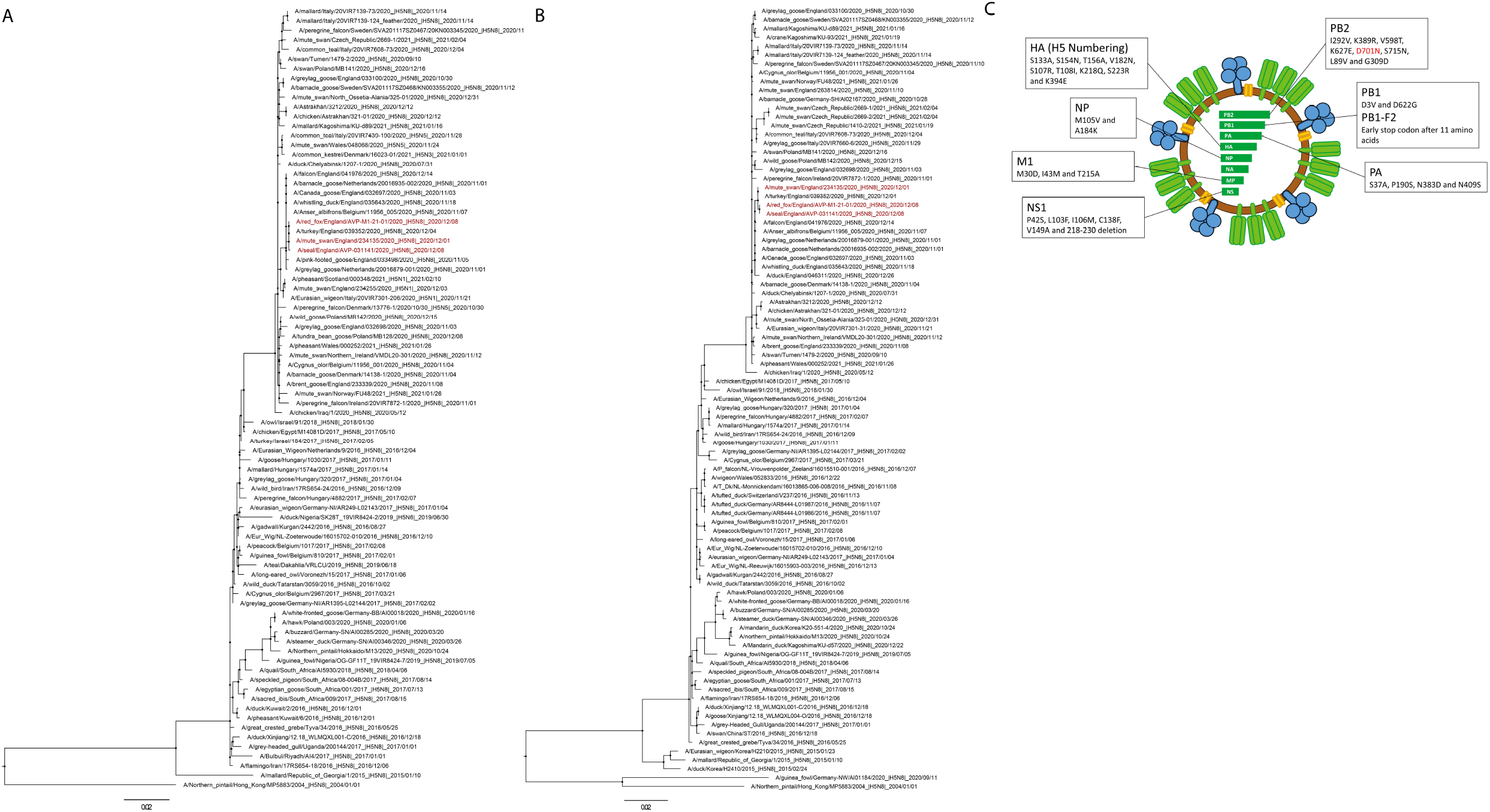
Genetic analyses of HA and NA from samples characterised from the disease event. A-Maximum-likelihood (ML) phylogenetic tree inferred from HA H5Nx segments predominantly from wild bird hosts. B-ML phylogenetic tree inferred from the NA segment of H5N8 viruses. All sequences generated in this study are coloured red; C-Viral schematic detailing amino acid changes present in the fox, seal and swan-derived viruses. Substitutions previously associated with altered phenotype in the literature are listed by gene segment. The D701N mutation detected only in the mammalian sequences is highlighted in red. Only H5 sequences from wild birds or poultry that were ancestral to the latest epizootic or contemporaneously detected in UK poultry outbreaks were included. Sequences were aligned with MAFFT v7.450 and trimmed to starting ATG and ending STOP codon using Aliview v1.26. Maximum-likelihood phylogenetic trees were generated with IQ-TREE v1.6.12 and branch supports were obtained with Shimodaira-Hasegawa-like approximate Likelihood-Ratio-Test (aLRT) with 1000 iterations. The final trees were visualised and annotated using FigTree v1.4.3 (https://github.com/rambaut/figtree) rooted through the outgroup A/Northern_pintail/Hong_Kong/MP5883/2004 and nodes placed in ascending order. The PDF is zoomable.

At the time of the outbreak, the rehabilitation centre fell within the Protection Zone (PZ) of a previously confirmed HPAI Infected Premises and, consequently, the off-movement of birds were restricted around the time of the disease episode. Restrictions were also applied specifically to the rehabilitation centre when suspicion of notifiable disease in the swans was reported. The cases investigated in this outbreak were limited to animals held within the isolation ward and over the subsequent month there were no further cases of unusual or unexplained clinical disease or mortality in mammals or birds at the centre. Following the disease event, but before diagnosis of H5N8 in the seals and fox, the centre was visited as part of surveillance activities within the PZ. Resident bird species were examined for evidence of clinical disease and 38 birds were sampled for virological testing (PCR). All birds tested negative.

## Discussion

Influenza A virus (H5N8) of avian origin was determined to be the cause of death in a red fox and the cause of seizures in a grey and several common seals present at a wildlife rehabilitation centre. This occurred roughly one week after the death of five swans due to HPAIV H5N8 housed in the same isolation unit. Genetic and epidemiologic investigations suggest that the swans were most likely the source of infection for the fox and seals, with transmission of virus by fomite transfer or aerosol spread.

The severity of disease and pathology in the seals and fox was unexpected. Outbreaks of disease associated with influenza A virus, with high mortality, have been described in Common seals associated with H10N7 [14-16], H7N7 [17] and H3N8 [18] viruses. In most of these cases, infection was limited to the respiratory tract. HPAI H5N8 clade 2.3.4.4 b has also recently been detected in Grey seals in Europe [19]. However, the infection of a fox as presented here is the first detection of this virus in terrestrial mammalian species and the first detection of this or any other influenza A viruses of avian origin in association with inflammation of the central nervous system in these species.

In 2016 and 2017, H5N8 was detected in lung tissue of two grey seals that were found dead, although pathology typical of influenza infection was not reported in either carcase [19]. Interestingly, infection with H5N8 in the seals in the presented case was associated with acute inflammation of the central nervous system; seemingly without viral replication or associated pathology in the respiratory tract, or detection in the nasal swabs collected prior to post mortem investigation, although there was evidence of viral nucleic acid in the lung of the single grey seal tested. The negative result on nasal swabs of the seals in this outbreak is noteworthy and may have implications for surveillance for influenza in this species.

Natural infection of terrestrial carnivores with influenza A subtypes, although rare, has been previously reported in species such as domestic and wild felids, mustelids and canids [20-22]. Experimental infection of red foxes was previously conducted using the HPAI H5N1 subtype by both intra-tracheal inoculation and through feeding virus-infected bird carcases (a proxy for the putative natural route of infection) [22]. This experimental H5N1 infection of foxes resulted in severe inflammation in the brain, lung and heart of foxes challenged intra-tracheally, despite the subjects showing no clinical signs. From the foxes fed virus infected carcases, only mild inflammation was seen, and it was restricted to the lungs. The study authors also refer in their discussion to detection of influenza A virus in foxes found dead in the field, although no further details were provided. The histopathological findings following the previously described intra-tracheal inoculated foxes are consistent with those observed in this natural fox infection and would support a respiratory or airborne infection route.

In the current study, evaluation of WGS data demonstrated that the virus present in the swans, the fox and the seals clustered together phylogenetically with minimal genetic differences observed. The WGS analysis did not enable the direction of infection to be determined, however, the epidemiological findings combined with the genetic data generated strongly suggests that the swans were the source of the infection for the fox and seals. The D701N amino acid substitution in the PB2 gene identified in all three sequences derived from mammalian species was absent from all avian sequences generated during the 2020/21 UK outbreak. This substitution has previously been associated with mammalian adaptation and increased replicative fitness in mammalian cells [23-27]. However, the substitution, in isolation, is not considered to be a factor that may result in increased avian-to-mammalian risk, with a combination of adaptive and compensatory changes being seen in human sequences and likely being required for efficient adaptation. Analysis of available H5 and H7 influenza sequences from human infections found that the D701N mutation had low prevalence and was therefore not a strong correlate for zoonotic infection. In conclusion, the assessment of the sequences derived from mammalian species, when compared against both avian influenza A viral sequences from the 2020/21 UK outbreak, and sequences derived from proposed human infection demonstrated no human risk over and above that already considered for the avian isolate.

A question remains as to why infection with a highly pathogenic H5N8 isolate of avian influenza produced such severe clinical disease in mammalian species in this event although contributing factors are likely to be multifactorial involving both viral and host elements. For example, a combination of co-morbidities, nutrition and physiological stress might have contributed to the development of disease in these animals. The health status of wild animals is difficult to establish, even when temporarily confined in wildlife rehabilitation centres. Several of the seals were admitted to the centre with lungworm infection. This is not uncommon in wild pinnipeds and, whilst the infections were severe enough to require treatment, several of the seals had been in the centre for at least a month and were reported to be responding well to treatment. The individual grey seal affected had been admitted as a neonate two weeks previously, and was likely to have been abandoned immediately after birth, probably without opportunity to suckle; therefore an immature immune system and reduced passive immunity could have been contributory factors in this seal. The fox was malnourished and suffering from mange, a common infection that causes severe morbidity and debility, and potentially increases susceptibility to other infections. Consideration was given to canine and phocine distemper viruses during the investigation, both as a potential cause of the neurological signs and encephalitis and as a cause of immunosuppression. However, this was excluded following negative results on PCR, WGS and IHC. There were no other pathologies or disease agents identified in the seals and fox at PME, although the range of testing was limited by autolysis of the carcases.

Malnutrition is common in young seals admitted to rehabilitation centres and was evident in the history provided for two of the common seals [28]. The centre was also feeding whole herring, which can present a risk of thiamine deficiency due to the presence of thiaminase in the fish. Fish from the centre subsequently analysed for thiamine returned markedly low levels and so it remains possible nutrition could have played a role in predisposing the seals to infection.

It is generally accepted that rehabilitation carries with it an unavoidable degree of stress to the animal, given handling, confinement and reduced opportunity to express natural behaviours. No definitive conclusions can be reached, but it could be hypothesised that the stress of confinement for rehabilitation may have stressed the affected animals, and impaired their immune response to viral or other, undisclosed infections.

In conclusion, influenza A virus of avian origin (subtype H5N8) was determined to be the cause of severe disease and mortality in seals and a fox held in a wildlife rehabilitation centre, along with swans which succumbed to the virus. All evidence suggested that the swans were the most likely source of infection for the fox and seals. Determining the cause of disease in the seals and fox was entirely reliant on collaboration between field veterinary services, pathologists and virologists and this case highlights the importance of wildlife disease surveillance. Whilst genetic analyses concluded there is no increased risk to humans of the H5N8 viruses in this outbreak, the investigation shows how these viruses may have unexpected and severe health risks for mammalian species. It is of note that there were no reports of illness in any staff associated with this event. However, such spill-over disease events in atypical host species constitute additional factors for veterinary authorities to consider during disease outbreaks and highlight the importance of wildlife disease surveillance utilising interdisciplinary and collaborative approaches.

## Acknowledgements

ACB, AMPB, EW, BCM, SMR, RH and IB were part funded by the UK Department for the Environment, Food and Rural A□airs (Defra) and the devolved Scottish and Welsh governments under grants SE2213, SV3400 and SV3006. The APHA Diseases of Wildlife Scheme is funded by Defra and the devolved Scottish and Welsh governments under grant ED1600. We thank the scientific and support staff of the Pathology and Virology departments of APHA Weybridge and of the Surveillance and Laboratory Services department of the APHA Veterinary Investigation Centres.

## References

1. Commission, E., Commission decision of 25 June 2010 on the implementation by Member States of surveillance programmes for avian influenza in poultry and wild birds (2010/367/EU) Annex II. Official Journal of the European Union, 2010: p. L. 166/22.

2. Núñez, A., et al., Highly Pathogenic Avian Influenza H5N8 Clade 2.3.4.4 Virus: Equivocal Pathogenicity and Implications for Surveillance Following Natural Infection in Breeder Ducks in the United Kingdom. Transbound Emerg Dis, 2016. 63(1): p. 5–9.

3. Commission, E., European Commision Decision 2006/437/EC of 4 August 2006 approving Diagnostic Manual for avian influenza as provided for in Council Directive 2005/94/EC of 4 August 2006 approving Diagnostic Manual for avian influenza as provided for in Council Directive 2005/94/EC. Official Journal of the European Union, 2006. 237: p. 1–27.

4. OIE, W.O.f.A.H. Terrestrial Manual: Avian Influenza (Infection with Avian Influenza Viruses) 2019 26 November 2019].

5. Seekings, A., et al., Highly pathogenic avian influenza virus H5N6 (clade 2.3.4.4b) has a preferable host tropism for waterfowl reflected in its inefficient transmission to terrestrial poultry. in press.

6. Marston, D.A., et al., Pan-lyssavirus Real Time RT-PCR for Rabies Diagnosis. J Vis Exp, 2019(149).

7. Gordon, C.H., et al., Canine distemper in endangered Ethiopian wolves. Emerg Infect Dis, 2015. 21(5): p. 824–32.

8. Fearnley, C., et al., The development of a real-time PCR to detect pathogenic Leptospira species in kidney tissue. Res Vet Sci, 2008. 85(1): p. 8–16.

9. Lewandowski, K., et al., Metagenomic Nanopore Sequencing of Influenza Virus Direct from Clinical Respiratory Samples. J Clin Microbiol, 2019. 58(1).

10. Li, H. and R. Durbin, Fast and accurate short read alignment with Burrows-Wheeler transform. Bioinformatics, 2009. 25(14): p. 1754–60.

11. Danecek, P., et al., Twelve years of SAMtools and BCFtools. Gigascience, 2021. 10(2).

12. Nurk, S., et al., Assembling single-cell genomes and mini-metagenomes from chimeric MDA products. J Comput Biol, 2013. 20(10): p. 714–37.

13. Pickett, B.E., et al., Virus pathogen database and analysis resource (ViPR): a comprehensive bioinformatics database and analysis resource for the coronavirus research community. Viruses, 2012. 4(11): p. 3209–26.

14. Bodewes, R., et al., Avian Influenza A(H10N7) virus-associated mass deaths among harbor seals. Emerging infectious diseases, 2015. 21(4): p. 720–722.

15. Krog, J.S., et al., Influenza A(H10N7) virus in dead harbor seals, Denmark. Emerg Infect Dis, 2015. 21(4): p. 684–7.

16. Zohari, S., et al., Avian influenza A(H10N7) virus involvement in mass mortality of harbour seals (Phoca vitulina) in Sweden, March through October 2014. Euro Surveill, 2014. 19(46).

17. Geraci, et al., Mass mortality of harbor seals: pneumonia associated with influenza A virus. Science, 1982. 215(4536): p. 1129–1131.

18. Anthony, S.J., et al., Emergence of fatal avian influenza in New England harbor seals. mBio, 2012. 3(4): p. e00166–12.

19. Shin, D.L., et al., Highly Pathogenic Avian Influenza A(H5N8) Virus in Gray Seals, Baltic Sea. Emerg Infect Dis, 2019. 25(12): p. 2295–2298.

20. Fiorentini, L., et al., Influenza A Pandemic (H1N1) 2009 Virus Outbreak in a Cat Colony in Italy. Zoonoses and Public Health, 2011. 58(8): p. 573–581.

21. Patterson, A.R., et al., Naturally occurring influenza infection in a ferret (Mustela putorius furo) colony. J Vet Diagn Invest, 2009. 21(4): p. 527–30.

22. Reperant, L.A., et al., Highly pathogenic avian influenza virus (H5N1) infection in red foxes fed infected bird carcasses. Emerg Infect Dis, 2008. 14(12): p. 1835–41.

23. Gao, Y., et al., Identification of amino acids in HA and PB2 critical for the transmission of H5N1 avian influenza viruses in a mammalian host. PLoS Pathog, 2009. 5(12): p. e1000709.

24. Le, Q.M., et al., Selection of H5N1 influenza virus PB2 during replication in humans. J Virol, 2009. 83(10): p. 5278–81.

25. Li, Z., et al., Molecular basis of replication of duck H5N1 influenza viruses in a mammalian mouse model. J Virol, 2005. 79(18): p. 12058–64.

26. Steel, J., et al., Transmission of influenza virus in a mammalian host is increased by PB2 amino acids 627K or 627E/701N. PLoS Pathog, 2009. 5(1): p. e1000252.

27. Taft, A.S., et al., Identification of mammalian-adapting mutations in the polymerase complex of an avian H5N1 influenza virus. Nat Commun, 2015. 6: p. 7491.

28. Colegrove, K.M., et al., Chapter 23 - Pinnipediae, in Pathology of Wildlife and Zoo Animals, K.A. Terio, D. McAloose, and J.S. Leger, Editors. 2018, Academic Press. p. 569–592.

